# Unraveling forensic timelines using molecular markers in *Phormia regina* maggots

**DOI:** 10.1101/2025.04.30.651557

**Authors:** Sheng-Hao Lin, Anthony J. Bellantuono, Kristian Lopez, Jeffrey D. Wells, Matthew DeGennaro

**Affiliations:** Department of Biological Sciences, Florida International University, Miami, FL 33199, USA; Biomolecular Sciences Institute, Florida International University, Miami, FL 33199, USA; Global Forensic and Justice Center, Florida International University, Miami, FL 33199, USA

## Abstract

In forensic entomology, estimating the time of death is critical and traditionally relies on changes in observable traits of carrion feeding insect larvae. Traits such as size, weight, and morphology can be used to predict the insect specimen age and help estimate the time since death. The blowfly *Phormia regina* is a key forensic insect, yet age estimation for older maggots in this species is particularly challenging due to the limited morphological changes in the late-stage larvae. To enhance precision, we employed transcriptomic profiling on blowfly maggots, aiming to identify genes as markers for time of death estimation. Our study characterized maggot development, reinforcing that weight and behavior cannot precisely determine age between 100 and 130 hours. We built a chromosomal scale annotated genome, establishing a reliable database for uncovering transcriptomic signatures during larval development. Applying differential gene expression analyses, weighted gene co-expression network analysis, and the generalized linear model, we identified nine candidate genes (*Y5078*, *Y5076*, *agt2, ech1, dhb4, asm, gabd, acohc, Ivd*) that delineate the age of otherwise indeterminate maggots. This research introduces a molecular approach to address a longstanding problem in forensic entomology and promises to increase precision in determining the time of death at a crime scene.

**AUTHOR SUMMARY:** In forensic entomology, estimating the time of death often depends on insect morphological changes. The blowfly *Phormia regina* is a key species, but aging older maggots in the third instar larval phase is difficult due to minimal changes in observable traits. To improve accuracy, we used transcriptomic profiling to identify genetic markers for time of death estimation. Our study showed that weight and behavior are not reliable indicators of age in maggots 100 to 130 hours old. We developed an annotated genome to identify transcriptomic signatures during larval development. Through various analyses, we identified nine genes (*Y5078*, *Y5076*, *agt2*, *ech1*, *dhb4*, *asm*, *gabd*, *acohc*, *Ivd*) that can determine the age of late third instar maggots. This molecular approach aims to enhance precision in forensic entomology.

## INTRODUCTION

Blowflies, members of the Calliphoridae family, are attracted to decaying organic matter like corpses by their keen sense of smell and sensitivity to decomposition cues [1]. Adult females typically deposit eggs on or near carrion because this is the larval food. These eggs hatch within hours to a few days, depending on environmental conditions, giving rise to maggots that progress through three larval stages before metamorphosing into adult flies. Knowledge of development rate under known environmental conditions, particularly temperature, is used to estimate the age of a specimen associated with a corpse. Since blowfly eggs are generally deposited on a cadaver after death, the age of the insect provides a critical minimum postmortem interval (PMI) estimate [2].

Estimating the postmortem interval (PMI) is a common step in a death investigation [2]. Although there have been methodological improvements to PMI estimation using many disciplines including microbiology [3], biochemistry [4], anthropology [5] and chemistry [6], the task is still very challenging because decomposition rate is a function of many variables, e.g. ambient temperature conditions and the effects of drugs in decomposing tissues [7]. There is a need for more precise methods to determine PMI that are less dependent on environmental variables [7].

The most common insect analyzed from a death scene is a blowfly [8]. The life cycle of a blowfly includes egg, three larval (maggot) instars, prepupa and pupa (both formed within the last larval exoskeleton, the puparium), and imago (adult) [9–11]. Unlike some other insects, blowfly maggots can become committed to metamorphosis very early during the final larval stage [12]. Therefore they can pupate and form a functional adult at a size much smaller than maximum, which appears to be an adaptation to the intense competition for food between both invertebrate and vertebrate animals that can occur on carrion [13]. In this investigation, *Phormia regina (P. regina)* was used as the study species. *P. regina* is common across North America except south Florida, often peaking in density during spring and fall [14–16].

The standard approach in forensic entomology for age estimation of immature maggots relies on the change of body length, weight, and instar [17]. During development larval size increases steadily [18], about midway through the third larval instar when larvae can stop feeding and wander in search of a location to pupate [19]. During most of the third larval instar the size of a larva does not change or may decrease slightly. As a result, neither instar nor size can be used to distinguish ages within about 50% of the larval life stage of *P. regina* and other carrion-feeding blowflies.

Age estimation methods based on changes in gene expression levels have been used in pupae of the blowflies *Calliphora vicina* and *Lucilia cuprina* [20,21] and the flesh fly, *Sarcophaga peregrina* [22]. Despite the need, age-related genetic markers of third instar blowfly maggots have not been identified. Some forensically important blowflies including *Cochliomyia macellaria* [23], *Lucilia sericata* [24] have shown that genetic markers can be associated with maggot age, but these studies do not provide detailed molecular information about third instar maggots over the critical developmental period where physical differences cannot distinguish maggot age.

In this study, we utilized transcriptome profiling to identify candidate genes to enhance the accuracy of blowfly maggot age estimation. We conducted a developmental study of *Phormia regina* maggots during their third larval instar. This blowfly species was chosen for its forensic importance across most of North America and Canada [25]. To support our investigation, we assembled a chromosomal-scale genome of *P. regina* as a robust reference for gene expression analysis and candidate gene identification. Additionally, we integrated Iso-Seq data to enhance genome annotation by providing full-length transcripts, which improve accuracy by enabling the direct detection of exon-intron boundaries [26]. We identified differentially expressed transcripts in maggots at 10-hour intervals during the third larval instar. We also found transcripts associated with maggots’ shift from feeding to wandering behavior. The candidate genes we have identified could be used by forensic entomologists to improve the accuracy of time of death determination.

## RESULTS

### Identification of key transitions during maggot development

Maggot developmental changes over time are used to determine when a blowfly laid her eggs on a corpse. To identify when it would be difficult to estimate the age of a maggot, we placed *P. regina* maggots of known age (∼24 h) on chicken liver (**Figure 1A**). To determine changes in phenotype with age, maggots were reared and sampled without replacement. Maggots were observed to be feeding if they remained on the liver substrate and were defined as wandering if they had ventured away from the cup, settling in the sawdust at the base of the rearing container (**Figure 1A**). We found a steady increase in larval weight and size until 100 hours old (h). From ages 110 h to 130 h no discernible changes in weight or size were detected. Highlighting the need of molecular determination of age for older maggots [27]. The first appearance of wandering maggots was observed at approximately 90 h, with this behavior becoming predominant by the 130 h mark (**Figure 1B and 1C**). Brown-Forsythe and Welch ANOVA tests indicated that the mean weight of the maggots increased between 70 and 100 h for both feeding and wandering maggots but there was no significant weight change between 110 and 130 h (**Figure 1B and 1C**). By applying the Mann-Whitney test to the weights based on behaviors, our results showed that the median weight of wandering maggots (66.10mg, n=307) is significantly higher than that of feeding maggots (34.55mg, n=507) (**Figure S1**). The transition from feeding to wandering also suggests a potential developmental shift in gene expression [28], indicating that investigating the gene expression profiles during larval development could yield valuable markers to enhance the precision of time since death estimation.

**Figure 1.**
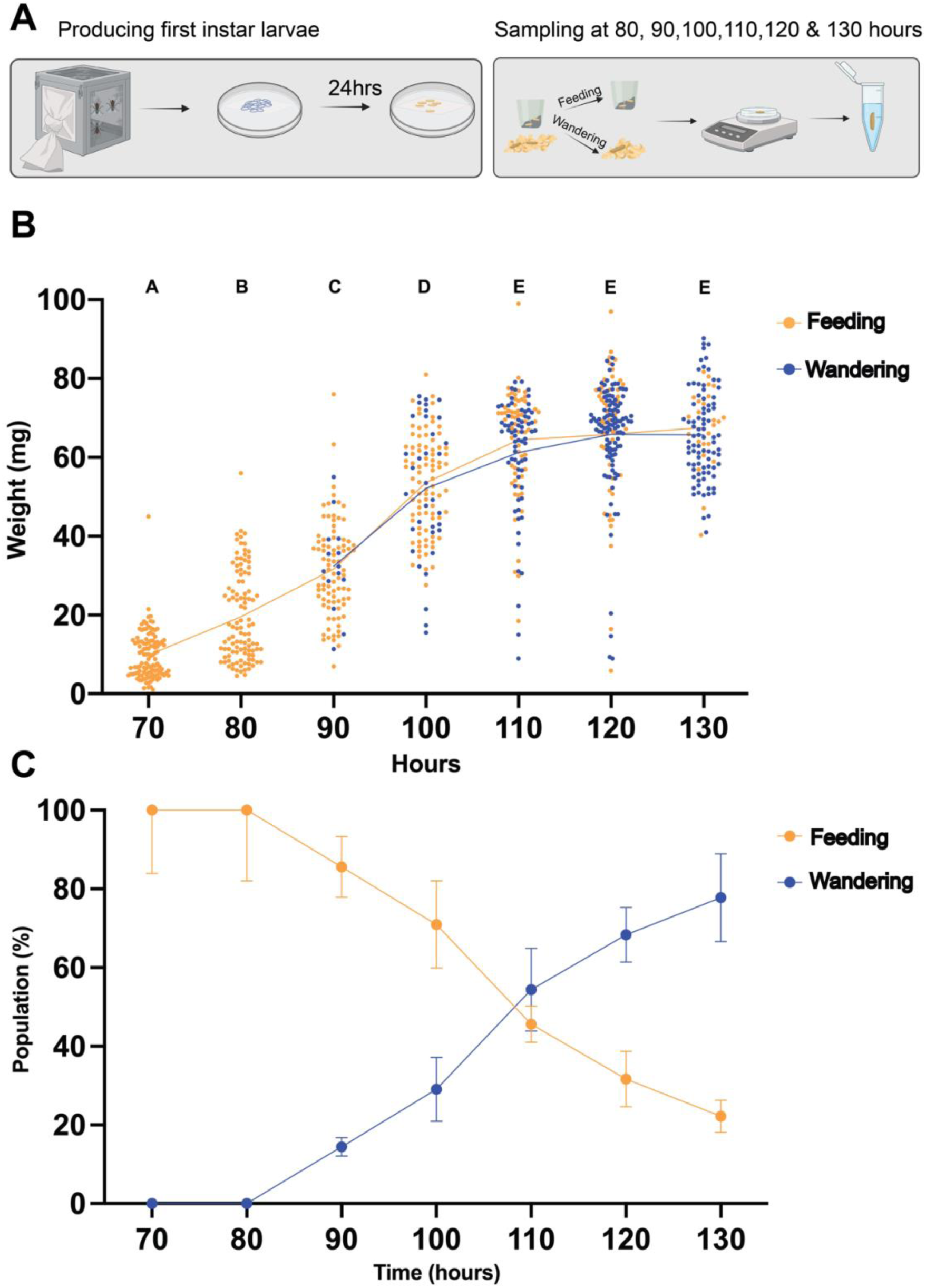
Tracking maggot weight and behavior from 70 hours old to 130 hours old. (A) Depiction of the sampling workflow where newly hatched maggots were transferred into a paper containing 30g chicken liver within a rearing box. The boxes were placed in a 27.5°C incubator. At each age cohort, maggots were visually scored for feeding or wandering behaviors and weighed. Created with BioRender.com. (B) Scatter plot of individually weighed maggots scored for feeding and wandering from 70 to 130 h. The straight line is the mean weight at each age cohort. The graph shows change in larval weight over four replicate experiments (n=823). Brown-Forsythe and Welch ANOVA tests were performed on the weight of aging cohorts (*P* < 0.0001). (C) The mean weight of feeding (yellow) and wandering (blue) maggots from 70 to 130 h. Population transition between feeding and wandering maggots occurred between 100∼120 h. The graphs showed the number of maggots scored as feeding or wandering (a) over four replicate experiments (n=823). The error bars indicate the standard error of each cohort.

### High quality *P. regina* genome assembly showed six pseudohaploid chromosomes

Prior to analyzing gene expression during larval development, we generated an updated, high-quality genome assembly of *P. regina* to serve as a reliable reference for transcript identification. Clean draft contigs were scaffolded using Hi-C data, and contact matrices were constructed to visualize chromatin architecture (**Figure 2A**; see **Materials and Methods**). Genome annotation was performed using the Funannotate pipeline, enabling the identification of protein-coding genes, including transcripts relevant to developmental stages. Assembly quality was assessed using a snail plot (**Figure 2B**), which showed an N50 of 8.2 Mb and a total genome size of 530 Mb. BUSCO analysis revealed a high completeness score with a low percentage of fragmented and missing orthologs, confirming the robustness of the assembly for downstream transcriptomic analyses. The plot also highlighted a low duplication rate among BUSCOs, suggesting minimal redundancy in the annotated genome. The Hi-C contact heatmap (**Figure 2C**) revealed six well-defined chromosomal territories, consistent with the known karyotype of *P. regina* [29], supporting our chromosomal-level scaffolding. These six pseudohaploid chromosomes were clearly delineated by blue boundaries, with red signals indicating high-frequency chromatin interactions within chromosomes. Comparative metrics (**Table 1**) show that the Hi-C-integrated assembly significantly improved genome contiguity, BUSCO completeness, and exon annotation, thus providing a robust genomic framework for transcript identification and expression profiling in *P. regina* [30].

**Figure 2.**
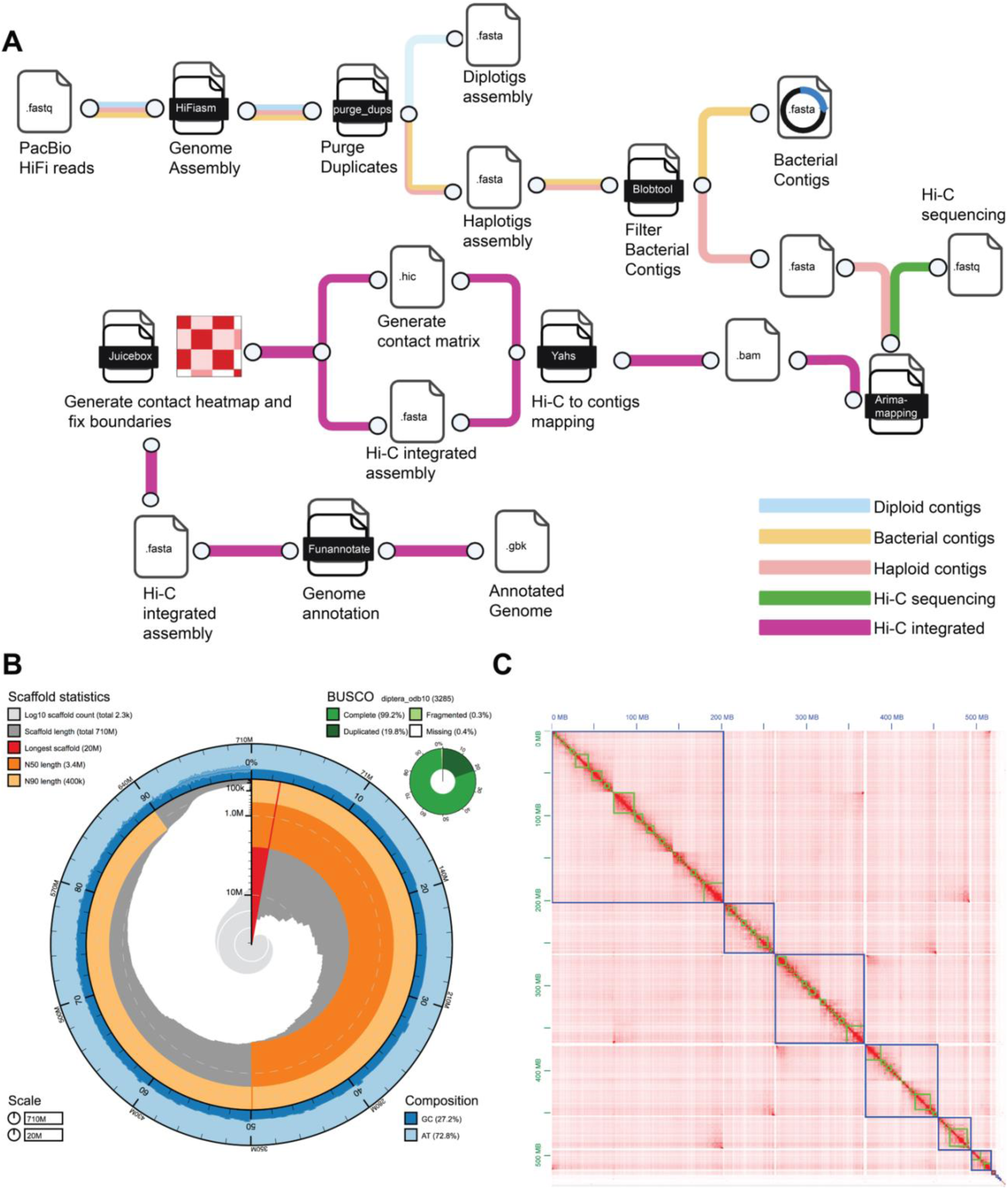
Chromosomal level genome assembly shows 6 pseudohaploid chromosomes in *Phormia regina*. (A) Genome assembly pipeline workflow. PacBio HiFi reads were assembled and duplicate sequences were purged. After the filtration of bacterial contaminant contigs from the assembly, the Hi-C sequences were mapped to the clean contigs for generating contact matrix and contact heat map. Created with BioRender.com. (B) Snail plots summarized the BUSCO, AT & GC content and scaffold statistics of *P. regina* genome assembly without proteobacteria contigs. Red segments represented the size of scaffold; dark and light orange segments indicated N50 and N90 values; central light gray indicated the cumulative scaffold counts under order of magnitude; blue and light blue indicate the proportion of AT & GC content. Upper right corner showed BUSCO completeness score against the Diptera database. (C) Hi-C contact matrix heat map of the chromatin interactions and scaffolding compartments. (Blue: Chromosomes, Green: scaffolds).

**Table 1.** Hi-C integrated *P. regina* genome assembly showed better quality than the published genome. The duplicates from the assembly were purged by HiFiasm and purge_dups. N50: median contig length. BUSCO: Benchmarking Universal Single-Copy Orthologs.

| Genome Assembly |  |  |
| --- | --- | --- |
| Quality | Male *P.regina* genome (Andere et al. 2016) | Hi-C integrated assembly (male) |
| No. of contigs | 1877 | 149 |
| N50(Megabase) | 7KB | 105.9MB |
| Total Length(Megabase) | 534MB | 534MB |
| Shortest contig(base) | 3 | 30,412 |
| BUSCO completeness(%) | 96.77 | 99.3 |
| Genome Annotation |  |  |
| Gene counts | 9490 | 28,518 |
| No. of genes hit BLAST | 8789 | 10,693 |
| No. of exons | 33,298 | 150,928 |

### *Agt2* and *Tsal* are associated with the transition between feeding and wandering behavior during the larval development

We identified 25 genes that displayed consistent upregulation and one downregulated gene when comparing age 90 h wandering to 90 h feeding behavior (**Figure 3A**). However, there were no differentially expressed genes while comparing feeding and wandering behaviors at other age cohorts except one upregulated gene at 120 h when wandering (**Figure 3B-F**). The distinct transcriptomic shift at 90 h corresponds to the initiation of the wandering phase—a developmental transition characterized by maggots leaving carrion and searching for a place to pupate (**Figure 1B**). The expression profiles indicated that most genes were downregulated during the wandering stage but upregulated during the feeding stages (**Figure S3A-B**).

**Figure 3.**
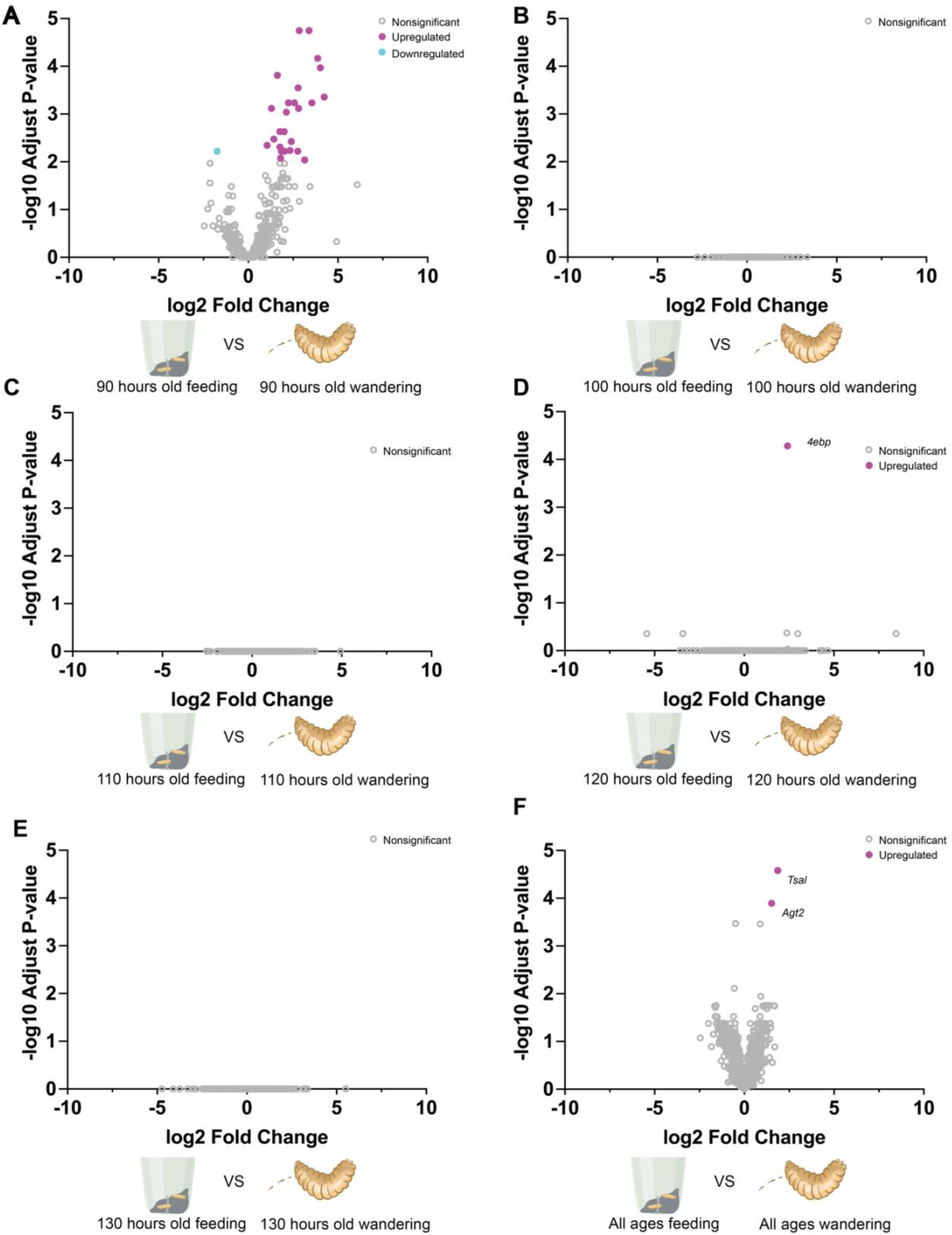
Age-dependent transcriptional changes associated with the transition from feeding to wandering in *P.regina* maggots. (A–F) Volcano plots showing differentially expressed genes (DEGs) between feeding and wandering larvae across different age cohorts: (A) 90 h, (B) 100 h, (C) 110 h, (D) 120 h, (E) 130 h, and (F) combined age groups. Differential expression was determined using DESeq2, applying a negative of log10 adjusted p value < 0.01 and log2 fold change > 1 as thresholds. Genes are color-coded: upregulated (magenta), downregulated (cyan), and nonsignificant (gray). Notable genes such as *4ebp*, *Agt2*, and *Tsal* are labeled in panels D and F. Statistical significance was assessed using the Wald test. NS denotes genes without significant differences in expression (∼10%).

### Dynamic Gene Expression Profiles During Blowfly Maggot Development Revealed by DEGs and WGCNA

When comparing the gene expression profiles of two conditions at older larval cohorts (90-130 h) to the 80 h cohort, volcano plots revealed the presence of twenty differentially expressed genes (DEGs), with a noticeable preponderance of downregulated genes in older larval cohorts (**Figure 4A**). Expanding our investigation to a comparison from other age groups (80 h, 100-130 h) to a 90 h cohort, volcano plots identified two upregulated and two downregulated DEGs (**Figure 4B**). In contrast, when we compared the gene expression profiles of samples at 100 h, 110 h, and 120 h to other age groups, the analysis did not reveal any significant DEGs (**Figure 4C-E**). We conducted a comprehensive analysis of Differentially Expressed Genes (DEGs) by comparing the earlier age cohorts (80 h-120 h) to a 130 h cohort. The resulting volcano plot showed a total of eighteen DEGs, the majority of which presented upregulated at early larval cohorts (**Figure 4F**). Using a heatmap of DEGs, we compared other age cohorts against the 80 h maggots; the heatmap emphasizes that most DEGs were upregulated between 80 to 90 h but downregulated in older maggots (**Figure S4A-C**). To ascertain the consistency of these gene expression patterns in the context of aging, we proceeded with Weighted Gene Co-expression Network Analysis (WGCNA), revealing a set of 168 genes characterized by co-expression patterns that either decreased or increased nearly synchronously over time (**Figure 5A**). Examination of turquoise, brown, and blue modules revealed different directional correlations in transcript expression across the age continuum (**Figure 5B**).

**Figure 4.**
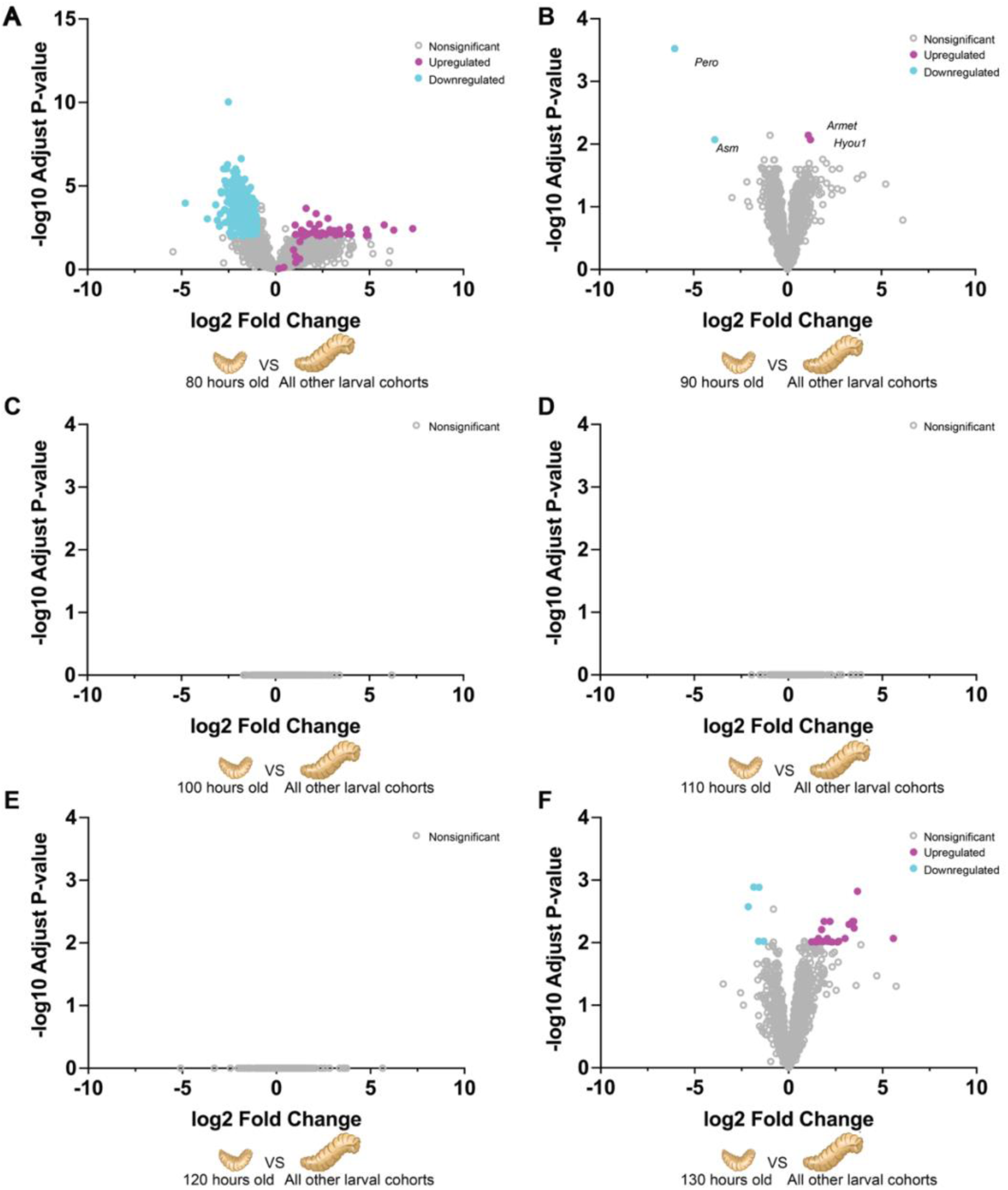
Age-specific gene expression signatures in *P. regina* maggots across developmental cohorts. (A–F) Volcano plots showing differentially expressed genes (DEGs) in each larval age group compared against all other age cohorts: (A) 80 h, (B) 90 h, (C) 100 h, (D) 110 h, (E) 120 h, and (F) 130 h. DEGs were identified using DESeq2, applying thresholds of log₂ fold change > 1 and adjusted *p*-value < 0.01 (–log₁₀ scale on the y-axis). Genes are color-coded as follows: upregulated (magenta), downregulated (cyan), and nonsignificant (gray). Notable differentially expressed genes such as *Pero*, *Asm*, *Armet*, and *Hyou1* are labeled in panel B. Statistical significance was assessed using the Wald test. NS denotes genes without significant differences in expression (∼10%).

**Figure 5.**
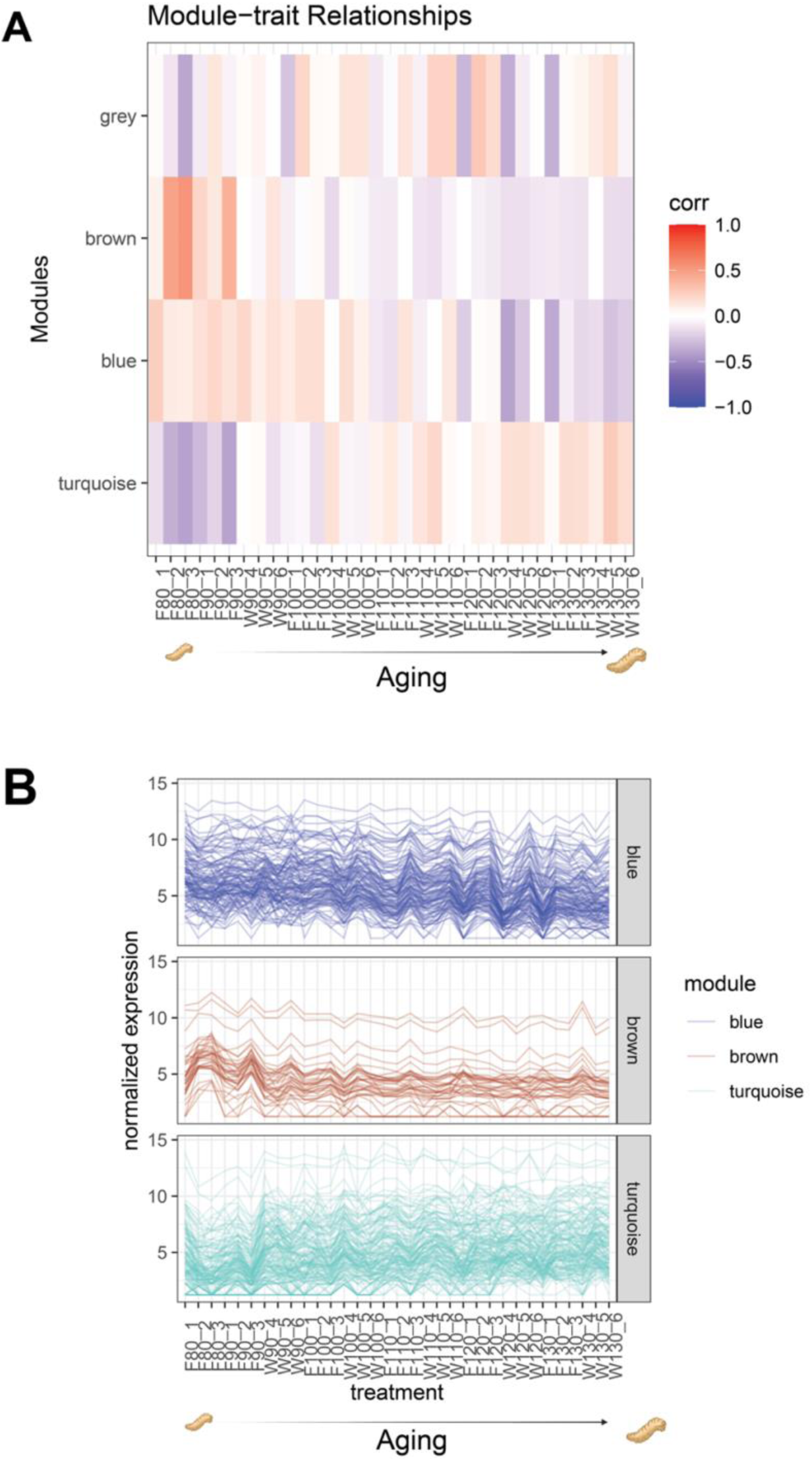
Weighted gene co-expression network analysis (WGCNA) reveals gene expression modules associated with developmental age and behavioral state in maggots. (A,) Heatmaps of module–trait correlations showing representation of normalized reads of sample clusters (column dendrograms) by developmental age. Color scale indicates the strength of correlation between module eigengenes. (B) Expression profiles of genes within selected WGCNA modules (blue, brown, turquoise) across samples, arranged by age. These modules exhibit coordinated gene expression patterns associated with developmental and behavioral transitions.

### GO Enrichment Analysis Identifies Molecular Pathways Associated with Blowfly Maggot Development

To gain insight into the biological relevance of the candidate genes identified from DEGs and WGCNA of wandering maggots, we conducted a statistical enrichment analysis of Gene Ontology (GO) terms. This analysis helped determine whether certain biological processes, cellular components, or molecular functions were overrepresented in each gene set compared to a reference set. We applied the DEGs list from the age cohorts to GO enrichment, revealing substantial involvement of differentially expressed genes at older larval cohorts (90-130 h), compared to the 80 h age cohort, in processes related to ribonucleoprotein, small nuclear ribonucleoprotein (snRNP) assembly, and ribosome components (**Figure 6A**). Conversely, the differentially expressed genes at 90 hours were associated with processes related to signal transduction (**Figure 6B**). Interestingly, the differentially expressed genes at 130 hours pointed to significant involvement in peroxidase activity (**Figure 6C**). Regarding behavioral transitions, we applied the gene lists from WGCNA and DEGs analysis to GO enrichment (**Figure 6D**). The results suggested that redox metabolism might influence wandering behavior prior to pupation, potentially due to anaerobic glycolysis in midgut metabolism [31]. Additionally, laccase activity was found in the epicuticle of wandering *L. cuprina* maggots, indicating a role for phenol oxidation in cuticular sclerotization while they are about to pupariate [32,33]. During the wandering stage, sterol transport and metabolism occur, likely contributing to the distribution of fat in adipose tissues for storage [34,35].

**Figure 6.**
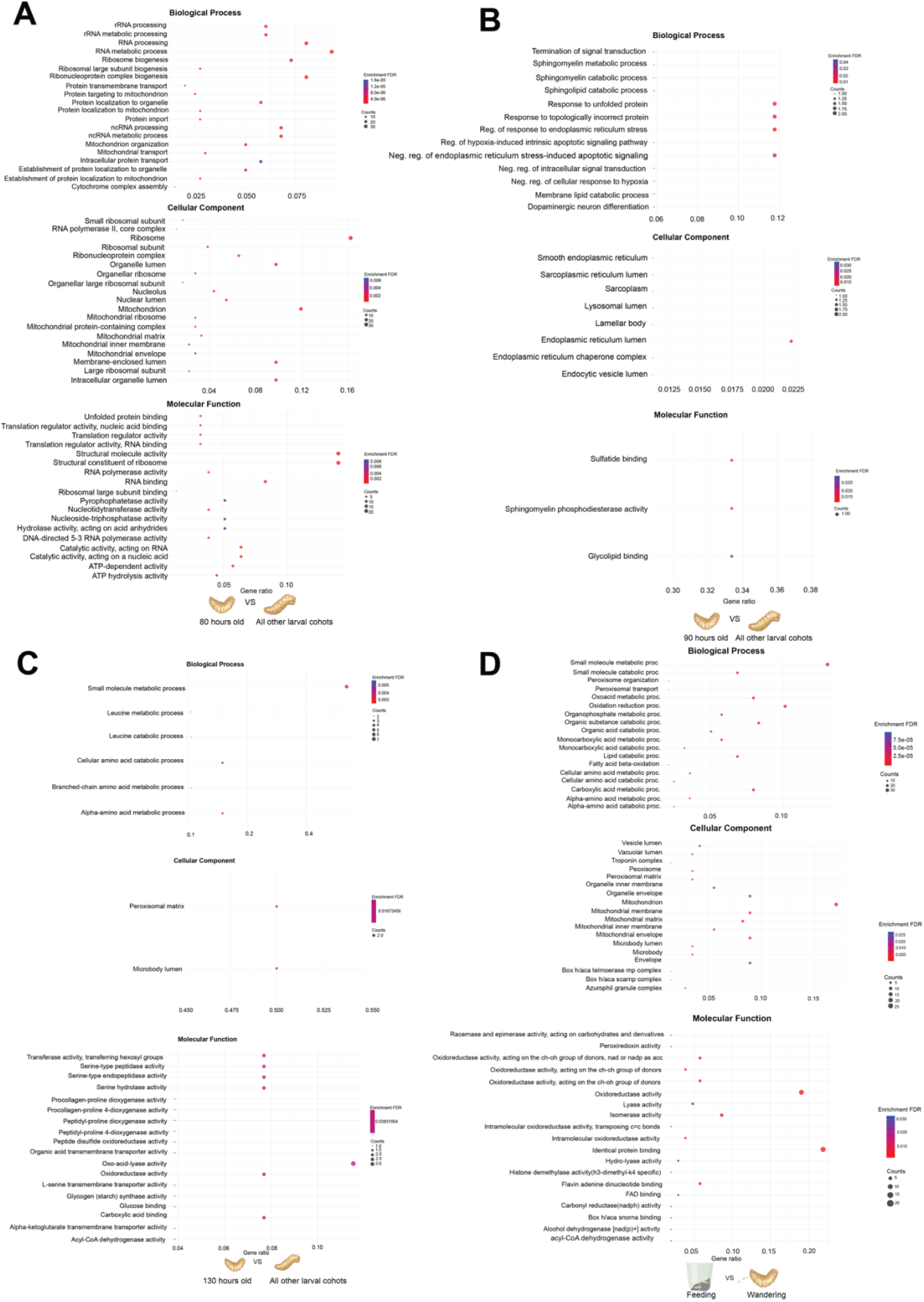
Gene ontology enrichment analysis highlights age- and behavior-associated metabolic pathways in maggots. This figure presents gene ontology (GO) enrichment analyses of differentially expressed genes (DEGs) across developmental timepoints (80 h, 90 h, 130 h) and behavioral states (feeding vs. wandering) in maggots. For each comparison, enriched GO terms in three categories—Biological Process, Cellular Component, and Molecular Function—were identified. Enrichment significance was determined using the Benjamini-Hochberg procedure with a false discovery rate (FDR) threshold of < 0.05. The Gene Ratio represents the proportion of DEGs associated with a specific GO term relative to the total number of input DEGs. Analyses reveal shifts in metabolic, cellular, and molecular pathways associated with age and behavior transitions. Created with BioRender.com.

### Nine genes associated with aging maggots are candidate molecular markers to improve forensic investigation

Unlike DEG analysis, which only provides information on significantly expressed genes, and WGCNA, which only offers correlations to certain clusters, utilizing general linear regression models (GLM) allows for the identification of useful markers that exhibit linear expression patterns over time, useful for accurate age determination of maggots in forensic investigations using methods such a quantitative or digital PCR. In our study, we applied the GLM to the normalized expression profiles of each gene over time, identifying a total of 58 upregulated genes and 44 downregulated genes that fit to this linear pattern (**Figure 7A-B**). We also identified 10 genes, characterized by unvarying TPM values, with *Dnlz* emerging as the gene with the highest TPM values (**Figure S7**). To narrow down our list of candidate genes, we utilized Venn diagrams to identify the overlaps among candidate genes identified through DEGs, GLM, and WGCNA analyses. Intriguingly, this analysis revealed nine genes that were identified in all three analyses, suggesting their potential utility in forensic applications (**Figure 8A**). Among these genes, *Y5078*, *Y5076*, *agt2, ech1, dhb4, asm* were upregulated; while *gabd, acohc, Ivd* were downregulated (**Figure 8B-C**).

**Figure 7.**
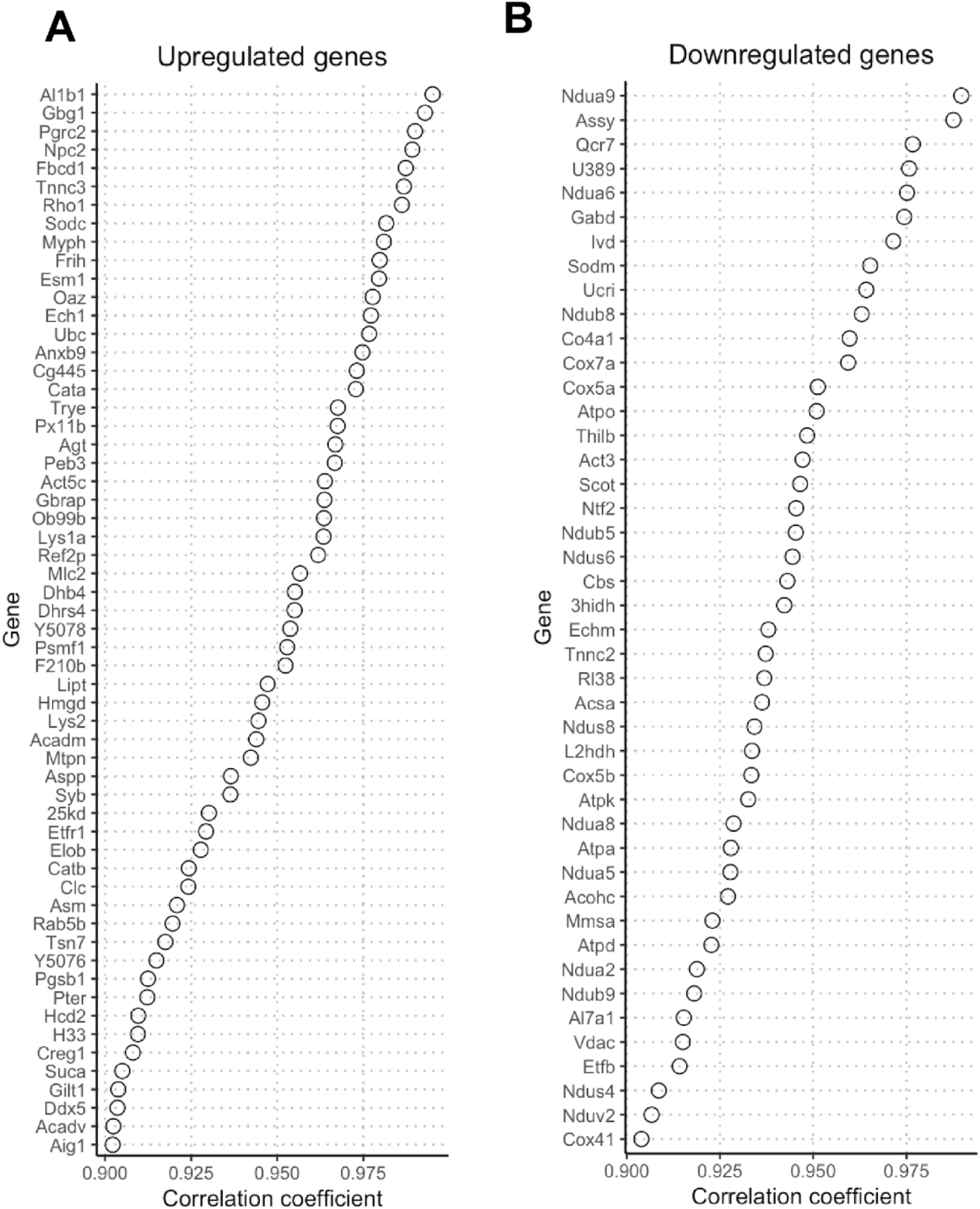
Age-associated candidate *P. regina* transcripts identified by linear regression analysis. Linear regression was used to identify transcripts with consistent changes in expression across developmental time points (80 h to 130 h). A total of 59 transcripts were classified as upregulated (A), showing increased expression over time with positive regression slopes (β₁ > 1) and strong model fit (*R²* ≥ 0.91). On the other hand, 45 transcripts were classified as downregulated (B), exhibiting decreased expression with negative slopes (β₁ < –1) and *R²* ≥ 0.91. Gene lists are ranked by correlation coefficient, and expression is reported in transcripts per million (TPM).

**Figure 8.**
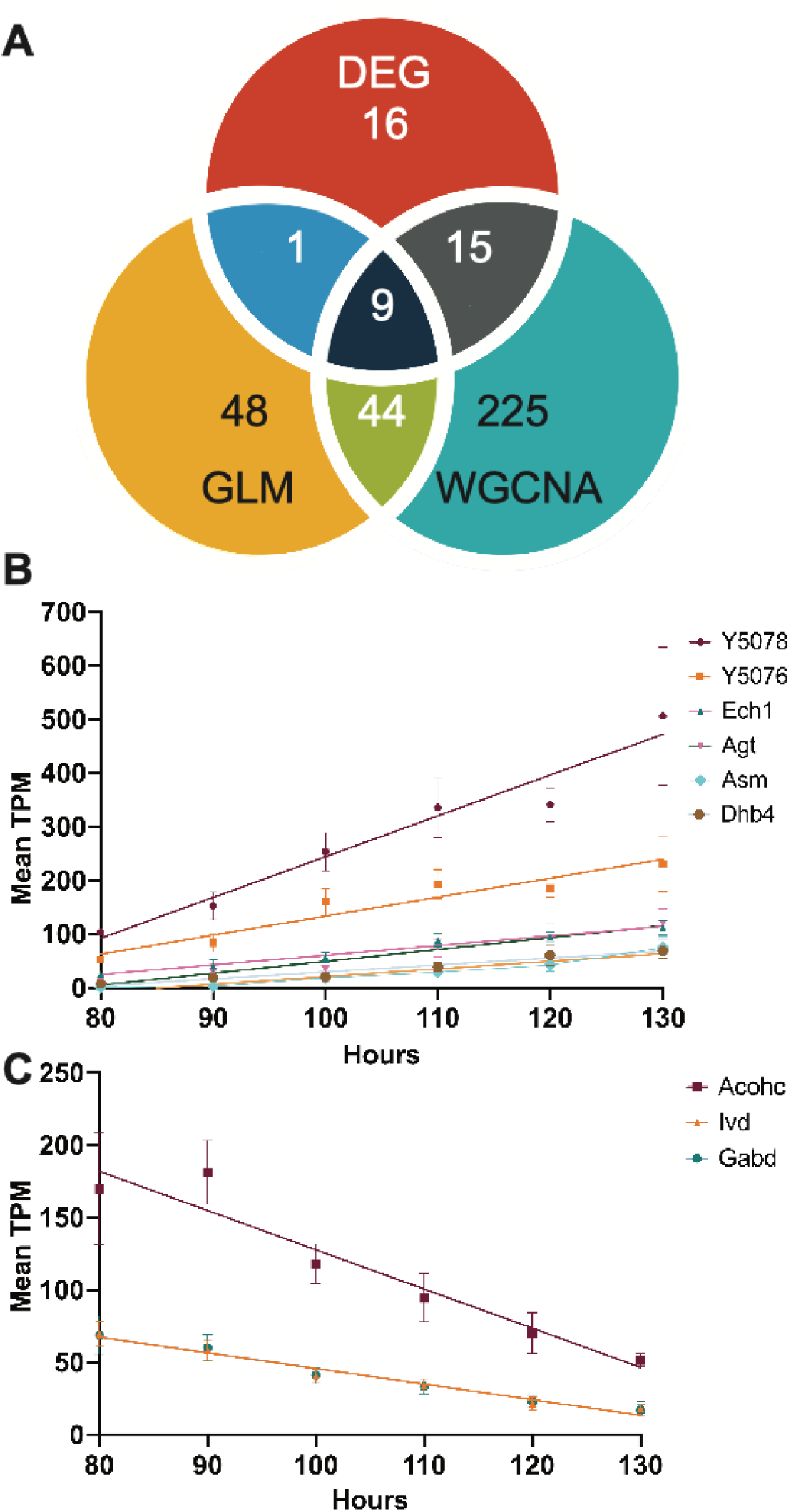
Nine candidate genes were among all three analyses. (A) Six candidate genes (*Y5078*, *Y5076*, *Agt, Asm, Ech1 dhb4*) appeared upregulated linear regression patterns. (B) Three candidate genes (*Gabd, Acohc, Ivd) appeared* downregulated linear regression patterns. (C) This Venn diagram displays a total of nine candidate genes that were consistently identified in all three analyses. GLM (General Linear Regression), WGCNA (Weighted Gene Co-Expression Network Analysis), and DE (Differentially Expressed Genes) analysis. The diagram illustrates the numbers of expressed transcripts associated with these three analytical approaches.

## DISCUSSION

Despite the practical and basic scientific value of the Calliphoridae, there has been relatively little genomic investigation of the group [36]. The high-quality, chromosomal-level genome assembly presented here represents a significant improvement over previous assemblies of blowflies [30]. This will help open up these forensically important insects to further genomic and genetic analyses. We used our updated genome to provide comprehensive gene models for our transcriptome analyses of maggot aging [37]. We identified 41 transcripts by DEG, 293 transcripts by WGCA, and 102 transcripts by GLM analyses. These genes are key candidates that are likely to improve age estimation of *P. regina* larvae. Out of these 358 transcripts, we highlight nine transcripts — *Y5078, Y5076, agt2*, *ech1*, *dhb4*, *asm*, *acohc, gabd* and *Ivd* — that were supported by all three methods of analysis.

We show that *Y5078* (Y5078_DROME uncharacterized protein from *Drosophila melanogaster*), *Y5076* (Y5076_DROME uncharacterized protein from *Drosophila melanogaster*), *agt2* (alanine glyoxylate aminotransferase), *ech1* (dienoyl-coenzyme A isomerase), *dhb4* (peroxisomal multifunctional enzyme type 2) and *asm* (sphingomyelin phosphodiesterase) are upregulated as maggots age. The genes *acohc* (cytoplasmic aconitate hydratase), *gabd* (succinate-semialdehyde dehydrogenase) and *Ivd* (Isovaleryl-CoA dehydrogenase) are downregulated as maggots age. These transcripts clearly delineate the aging of maggots from 80 h to 130 h making them suitable markers for age estimation. In addition, our results provide new insights into the regulatory network underpinning homeostasis and metabolism during third instar maggot development. For example, the upregulated marker genes have been associated with cholesterol export, energy metabolism, fatty acid metabolism, and detoxification [38–42]. On the other hand, downregulated marker genes participate in regulation of iron homeostasis and GABA degradation pathway as maggots age [43,44].

Our study identified 27 transcripts from DEGs and 293 transcripts from WGCNA associated with feeding and wandering behavior. Notably, the presence of DEGs at 90 hours, consistent with the presence of wandering maggots at 90 hours, suggests a strong connection between these genes and the behavioral transition. Two genes, *Agt2* (alanine glyoxylate aminotransferase) and *Tsal* (L-threonine ammonia-lyase), were significant from the two analyses, indicating a potential role for amino acid homeostasis during transition from feeding to wandering. [45,46].

While larval size is often used as an age indicator, it remains an imperfect metric due to substantial variation in size at a given age, particularly during later larval development [47,48]. Similarly, as seen in these results the age of transition from feeding to wandering is also highly variable, making that milestone of limited use for estimating age (**Figure 1B**.). Gene expression provides a more reliable framework for age estimation because conventional age estimation by morphological traits can be more easily influenced by the variability in environmental conditions like diet and

temperature, which significantly affect body size [49]. By combining enhanced transcriptional information, our study identifies candidate genes that not only contribute to a better understanding of dipteran development and behavior during the third instar larval stage but also have potential applications in forensic investigations for estimating time since death. Further field studies are needed to validate the performance of these candidate genes in *P. regina* and to facilitate the development of genetic marker kits that can ultimately improve precision in time-of-death estimation.

## MATERIAL AND METHODS

### Identifying the transition between feeding and wandering behavior

We maintained a laboratory colony of an inbred line of *Phormia regina* originally from Lincoln, Nebraska [50]. Developmental rate data were available [50]. All rearing containers were maintained at 27.5± 0.5 °C. under 16:8 light:dark cycle in one SMY04-1 DigiTherm CirKinetics Incubator (TriTech Research, Inc., Los Angeles, California, USA). The incubator was equipped with uniform lighting, additional fans, and a port for thermometer access.

Each equal-age cohort was reared within a plastic 72 x72×100 mm insect breeding box (Gyeonggi-do, Korea). Each box contained 0.5cm sawdust and a 30g of fresh chicken liver in a suspended 4 oz paper cup. Eggs were obtained on a paper towel soaked with chicken liver blood placed in a cage of *P. regina* adults for 30 minutes before removal. The removal time was considered age zero for those insects. About 500-1000 newly deposited eggs were placed on wet paper in a covered petri dish. The eggs were kept at high humidity over an open water container at 27.5± 0.5 °C. After 24 hours, fifteen newly hatched maggots were transferred into each aliquot of fresh liver inside the rearing box (See **Fig 1A**). All of the boxes were reared at the same 27.5± 0.5°C incubator. For each of four trials, 28 boxes with maggots were set up. During each sampling time, four rearing boxes were randomly removed from the incubators. We distinguished a maggot as feeding if it was still on the liver and wandering if it was in the sawdust. Sampling involved removal of an entire cohort at a preselected age. Sampled maggot was individually placed in a 1.5mL microcentrifuge tube with 1mL of Thermofisher RNAlater (Invitrogen, MA, USA) at 4 °C. Each tube was labeled with numbers for RNA analysis. After 24 hours, the RNAlater was removed from the tube and the insect was stored at −80 °C.

### RNA & DNA sequencing

Over three independent rearing cohorts, individual maggots were selected based on their age, behavior, and average weight relative to other members of the same cohort. Maggots exhibiting feeding behavior (F) or wandering behavior (W) were collected at 10-hour intervals from 80 to 130 hours post-hatching. The specimens were homogenized using a pestle with 200 μl of ambion TRIzol reagent (MA, USA). RNA was extracted from using a QIAGEN RNeasy Plus extraction kit (MA, USA). The concentration of the RNA samples was assessed with an Invitrogen Qubit Fluorometer using a Qubit RNA HS Assay kit (MA, USA). The integrity of the extracted RNA was evaluated with a Bioanalyzer using an Agilent RNA 6000 Pico chip. A total of 33 RNA samples from maggots of known ages were sent to BGI (CA, USA) for RNA sequencing, and 2 RNA samples from adult flies following the same RNA extraction as previously described were sent to Genewiz for RNA sequencing (NJ, USA).

A male virgin adult blowfly was sent to PacBio for sequencing using the PacBio Sequel II platform to generate a HiFi read library. Additionally, another male virgin adult blowfly was sent to CD Genomics, where Hi-C libraries were quantified and sequenced on the Illumina NovaSeq platform (San Diego, CA, USA). See **Supplemental Table 1** for details on short-read and long-read libraries.

### Iso-seq mRNA analysis of *P. regina*

A single virgin adult male, a single virgin adult female, a 110 h feeding third instar maggot, and a 110 h wandering maggot were sent to DNA Sequencing Center/Brigham Young University for Iso-seq. Iso-Sequencing was performed on the PacBio Sequel II platform. Consensus sequences were generated from the SMRTBell libraries and collapsed into isoforms by Isoseq v3.0 [51]. The long reads libraries were used as evidence for genome annotation.

### Build a new *P. regina* genome assembly

We performed *de novo* assembly using HiFiasm v0.16.1 on a HiFi long-read library from a single adult male blowfly (See **RNA & DNA sequencing**), adopting purge mode to generate both haplotype and diplotype assemblies of *Phormia regina* [52]. Duplicate contigs were further eliminated by purge_dups v1.2.6 [53]. The contaminated contigs were identified by NCBI BLAST against a publicly available nucleotide database and filtered by Blobtool v2.6.4[54]. The repeat element boundaries and repeat database was *de novo* assembly by the clean contigs using RepeatModeler v2.0.2 [55–63]. Hi-C sequencing libraries from a male adult blowfly (See **RNA & DNA sequencing**), were first mapped to the clean contigs by Arima mapping pipeline and the .bam file was used to perform chromosomal integrated assembly by yahs v1.1 [64]. A Hi-C heat map was generated by juicebox v2.20.00 [65]. The curation and chromosomal boundaries were manually edited following the Genome Assembly Cookbook [66]. The chromosomal genome was annotated using the Funannotate v1.8.13 annotation pipeline, with Iso-seq libraries providing support for evidence-based gene prediction during the annotation process. (See **Iso-seq mRNA analysis of *P. regina***) [67–72].

### Gene expression analysis

The quality of RNA seq libraries previously described (See **RNA & DNA sequencing**) were initially assessed by FastQC v0.11.9 [73]. The adapter sequences from the short reads were trimmed by Trimmomatic v0.39 [74]. The clean short read libraries were mapped to the newly annotated genome (See **Build a new *P. regina* genome assembly**) using Salmon v1.8.0 for quantification [75]. Count matrices were obtained by tximport 1.34.0 [76]. Differential expression analysis was performed by DeSeq2 1.34.0 Bioconductor [77]. The gene names were manually curated based on the results from the Blastn against orthologous UniProt v2023_02. Pairwise comparison was performed among all treatments to identify differentially expressed transcripts (> 2-fold change, *a* < 0.01 FDR) using DEBrowser V3.18[78]. Gene co-expression analysis was produced by DESeq2 v1.34.0 followed by tidyverse v2.0.0, magrittr v2.0.3, and WGCNA v1.72.1 analysis on R v2022.12.0+353 [79,80]. Gene ontology enrichment analysis was performed by using ShinyGO v0.76 under FDR-corrected cut off *a* < 0.05 [81].

### Finding candidate genes for predicting larval age

Raw reads from the development of aging maggot as previously described were normalized to TPM values by tximport v1.34.0 [76]. Linear regression model was applied to define the housekeeping genes (*s* <-2, all values greater than 0), upregulated genes (b*_1_* > 1 and *R^2^*: 0.9∼1) and downregulated genes ( *b_1_* < −1 and *R^2^*: 0.9∼1) from the mean TPM values across each age cohort; mean TPM values from the linear regression model were calculated over six replicates (including three feeding replicates and three wandering replicates) for age cohorts from 90 to 130 hours, and the 80-hour time point includes only three feeding replicates (See **Supplemental Table 1**). The correlation plots were created using ggplot2 [82] The Venn diagram list was created by the VennDiagram v1.7.3 package from RStudio v0.15.0 and drawn by Procreate v5.3.7 from Savage Interactive Pty Ltd [83,84]. Linear regression figures were created by Prism10 v0.2.2 from GraphPad software.

## DATA AVAILABILITY

The RNA and DNA sequencing data files are available for download on NCBI Sequence Read Archive with Bioproject ID PRJNA990781. The annotated genome is available for download with Bioproject ID PRJNA1085338. All the other underlying data for the manuscript will be made available via dryad data repository.

## ACKNOWLEDGMENTS

Dr. Andre Luis Costa-da-Silva (Florida International University, FIU) provided invaluable guidance on the molecular work. Dr. Philip Stoddard (FIU) designed and 3D-printed equipment for our behavioral assay. Brian Haas (The Broad Institute) helped us troubleshoot and refine the PASA aligner pipeline. Mali O. Jones from contributed artwork for the figures and Venn diagrams featured in this paper. Our laboratory colony of *P. regina* was established with specimens from Dr. Amada Roe (College of Saint Mary). We thank Dr. Amber MacInnis for sharing her expertise in rearing blowflies. S.L. was supported by NIH grant 5T32GM132054-05 and the Department of Biological Sciences at Florida International University. The work was supported by National Institute of Justice grant 2019-DU-BX-0013 to J.D.W. and M.D. We extend our gratitude to the Florida International University Instructional Research & Computing Center for providing the computational resources.

## AUTHOR CONTRIBUTIONS

**Conceptualization:** Sheng-Hao Lin, Jeffrey D. Wells, Matthew DeGennaro.

**Data curation:** Sheng-Hao Lin, Kristian Lopez, Anthony J. Bellantuono,

**Formal analysis:** Sheng-Hao Lin, Anthony J. Bellantuono.

**Funding acquisition:** Jeffrey D. Wells, Matthew DeGennaro.

**Investigation:** Sheng-Hao Lin, Kristian Lopez, Anthony J. Bellantuono.

**Project administration:** Jeffrey D. Wells, Matthew DeGennaro.

**Supervision:** Matthew DeGennaro.

**Visualization:** Sheng-Hao Lin.

**Writing – original draft:** Sheng-Hao Lin.

**Writing – review & editing:** Sheng-Hao Lin, Anthony J. Bellantuono, Jeffrey D. Wells, Matthew DeGennaro.

## SUPPLEMENTARY FIGURES

**Figure S1, related to Figure 1.**
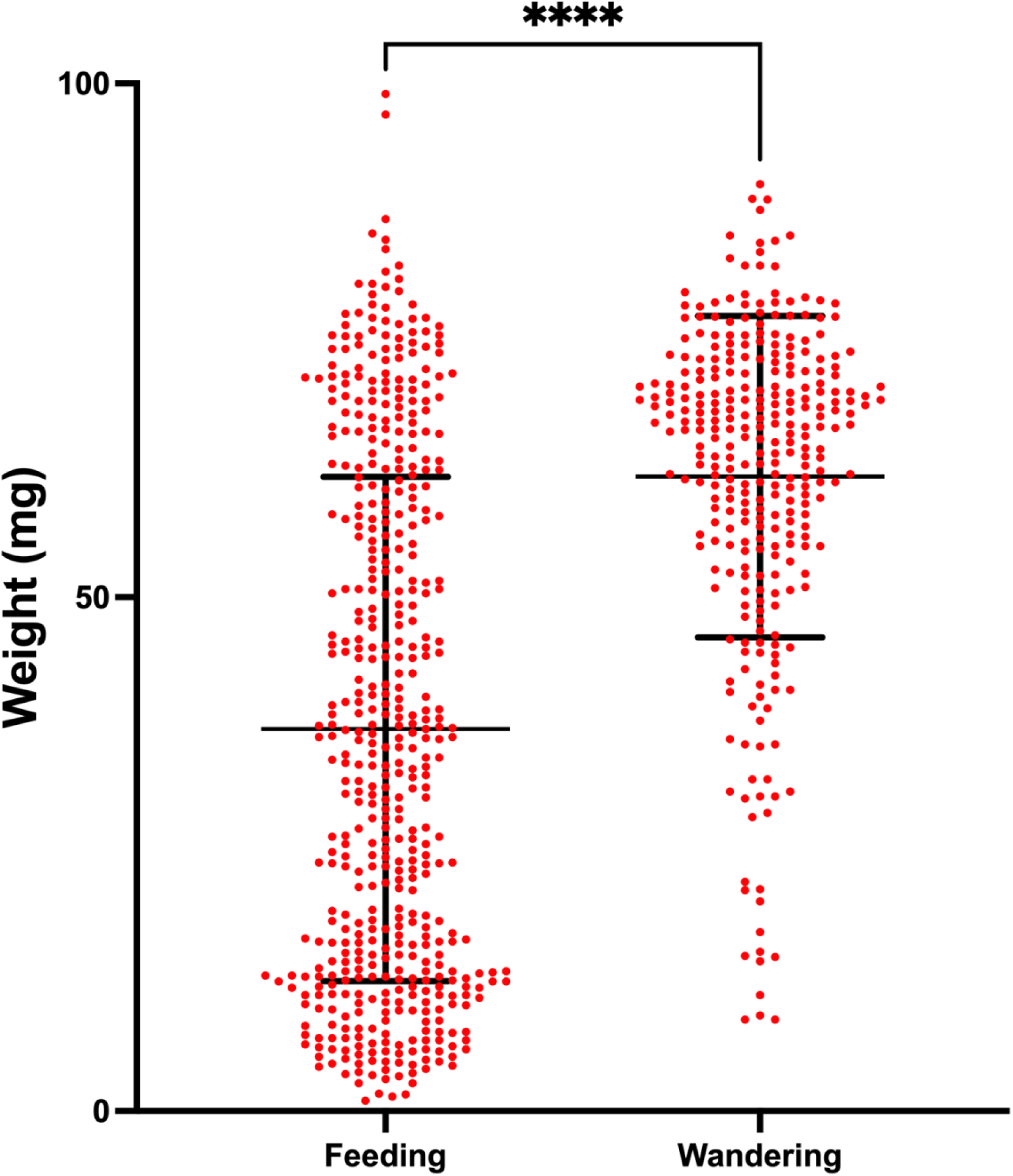
Median weight of wandering maggots is significantly higher than the median weight of feeding maggots. The figure presents findings of comparing the weights of feeding and wandering maggots across ten age cohorts spanning from 80 to 130 h. Maggots were categorized into feeding or wandering groups based on their behavior. The median weight of wandering maggots significantly exceeded that of feeding maggots according to the Mann-Whitney test (*p* < 0.0001). Each maggot’s weight is depicted by a dot, with the center horizontal line of each box indicating the median weight. Additionally, error bars depict the standard error.

**Figure S3, related to Figure 3.**
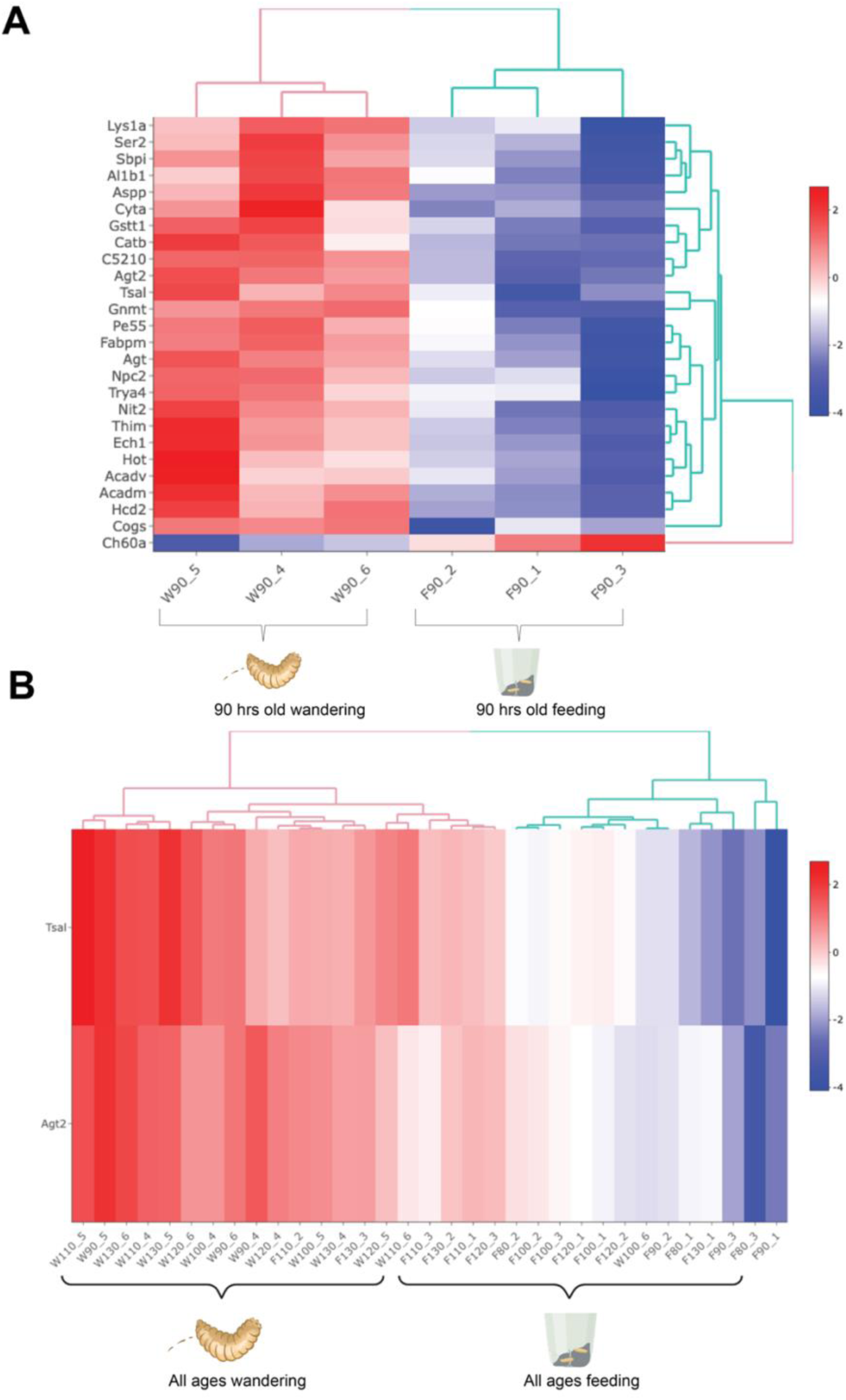
*Agt2* and *Tsal* were differentially expressed genes in feeding and wandering behavior during larval development. The heat map showed differentiated expressed transcripts over aging maggots sorted into feeding (F) and wandering (W) at 90 h (A) and all the ages (B) behaviors from 80 h to 130 h. The figure showed the clusters of relationships (y axis) among the transcripts to the left and treatment on the x axis (numbers: age from 80 to 130 h). The heat map was generated by DESEq2 based on the Spearman correlation (fold change > 2, FDR < 0.01). Created with BioRender.com.

**Figure S4, related to Figure 4.**
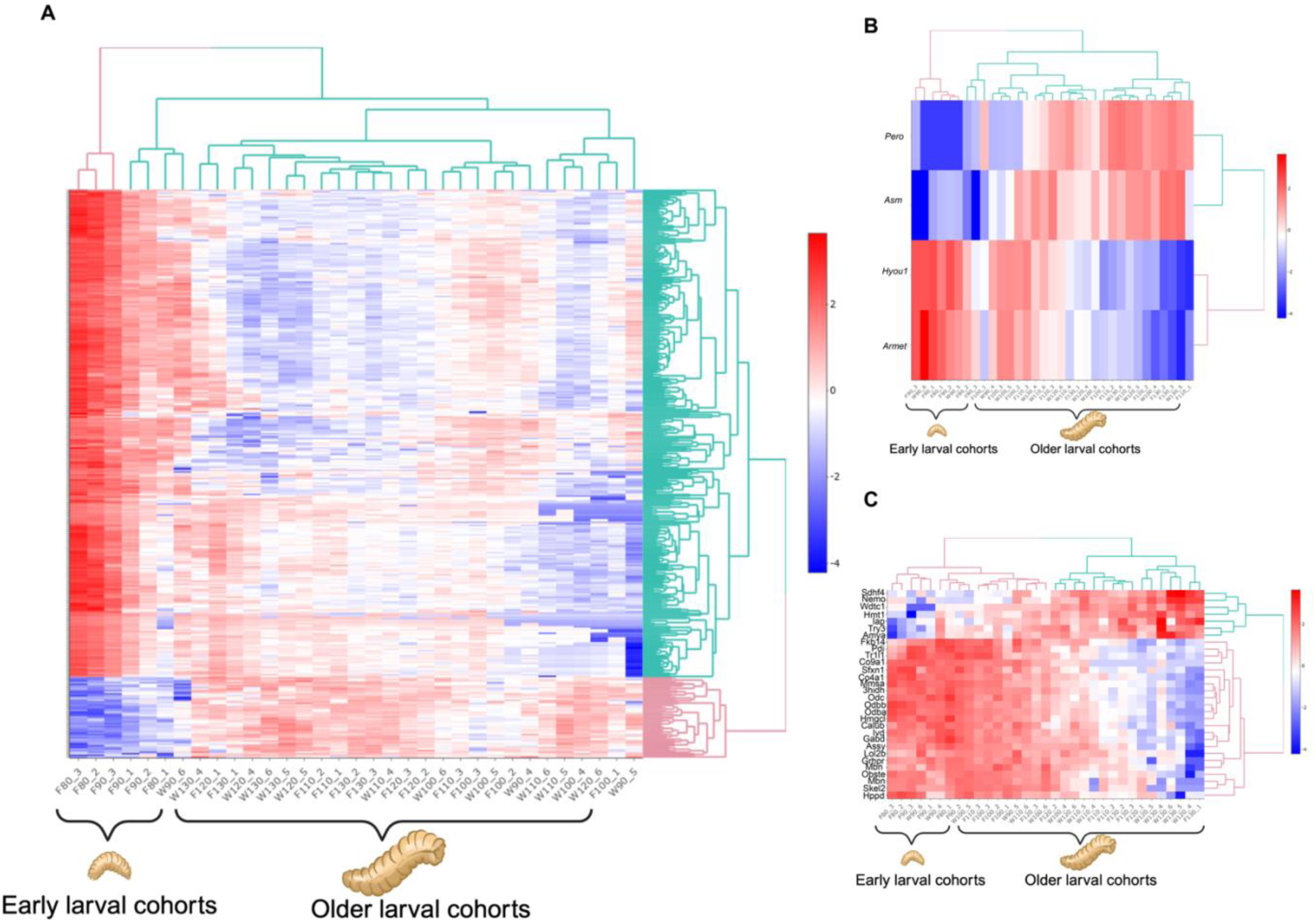
Heatmaps reveal distinct expression patterns of transcripts when comparing early and older aging cohorts. Three pairwise comparisons were conducted: (A) 80 h versus other aging cohorts, (B) 90 h old versus other aging cohorts, and (C) 130 h versus other aging cohorts, resulting in heat maps displaying distinct expression patterns of transcripts between early and older aging cohorts. Clusters of relationships among transcripts are presented on the right, with the treatment depicted on the x-axis (feeding (F), wandering (W), numbers: age from 80 to 130 h). The heat map, generated by DESEq2 using Spearman correlation, focuses on transcripts with fold change >2 and FDR < 0.01.

**Figure S8, related to Figure 8.**
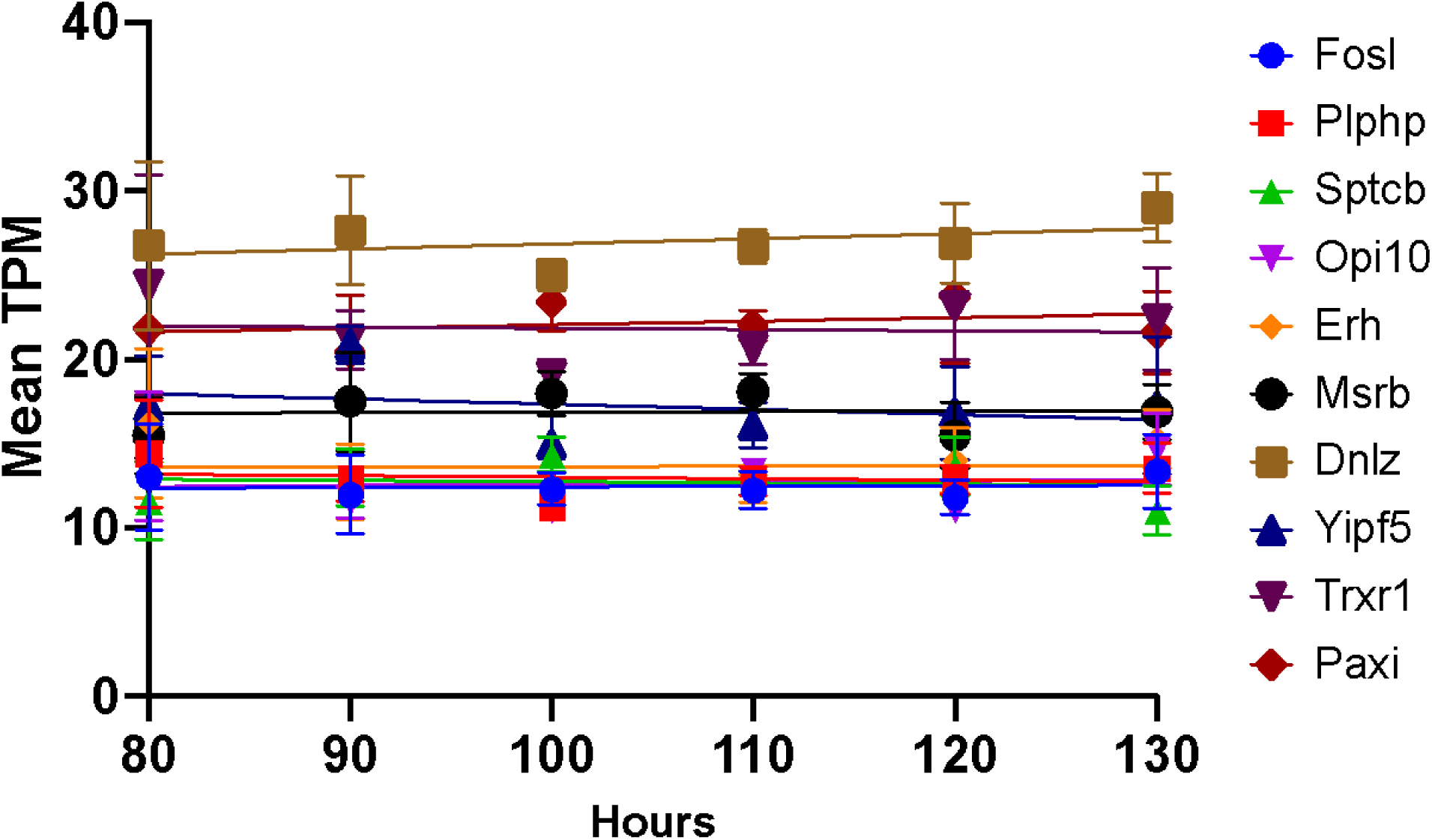
Linear regression analysis showed the genes whose expression did not vary over developmental time. Linear regression model showed 10 housekeeping genes expressed over time (80 h ∼130 h) at transcripts per millions (TPM) values (*s* <-2, all values greater than 0).

